# REM sleep promotes bidirectional plasticity in developing visual cortex *in vivo*

**DOI:** 10.1101/2022.01.21.477249

**Authors:** Leslie Renouard, Christopher Hayworth, Michael Rempe, Will Clegern, Jonathan Wisor, Marcos G. Frank

## Abstract

Sleep is required for the full expression of plasticity during the visual critical period (CP). However, the precise role of rapid-eye-movement (REM) sleep in this process is undetermined. Previous studies in rodents indicate that REM sleep weakens cortical circuits following MD, but this has been explored in only one class of cortical neuron (layer 5 apical dendrites). We investigated the role of REM sleep in ocular dominance plasticity (ODP) in layer 2/3 neurons using 2-photon calcium imaging in awake CP mice. In contrast to findings in layer 5 neurons, we find that REM sleep promotes changes consistent with synaptic strengthening and weakening. This supports recent suggestions that the effects of sleep on plasticity are highly dependent upon the type of circuit and preceding waking experience.

## Introduction

Sleep enhances a classic model of developmental plasticity *in vivo* (ocular dominance plasticity: ODP) ^1^. ODP is induced by monocular deprivation (MD) during a critical period (CP) of development, resulting in a shift of cortical response to the non-deprived eye (NDE). ODP is considered physiological as it involves adaptive changes in response to sensory input and normally governs the proper development of binocular vision. ODP is also considered a canonical form of plasticity as many of the underlying mechanisms discovered in this system govern diverse forms of plasticity elsewhere in the brain. Therefore, what we learn about sleep in ODP may reveal general rules about sleep function ^2–4^.

The precise role of rapid-eye-movement (REM) sleep in ODP is unknown. Previous studies indicate that REM sleep is necessary for ODP, as REM sleep deprivation (RSD) in CP cats and mice reduce shifts in cortical response to the NDE ^5, 6^. The types of plasticity governing this shift are unclear. In cats, sleep strengthens responses to the NDE, but the specific role of REM sleep in this process was not investigated ^5, 7^. In mice, REM sleep weakens responses to the deprived eye (DE), but this analysis was restricted to one type of neuron (apical dendrites from layer 5 neurons) ^6^. Considering that excitatory neurons in cortical layers 2/3 are among the first to exhibit ODP^8^, this raises the possibility of a more heterogenous response.

To determine the role of REM sleep more precisely in ODP, we used a combination of 2-photon fluorescence microscopy and polysomnography in CP mice expressing GCaMP6s in layer 2/3 CaMKII+ neurons (Figure 1). We find that short periods of MD and sleep lead to more complex changes than previously reported, including a gain of response to the NDE. This supports the idea that sleep promotes bidirectional changes in plasticity depending on the cell and circuit ^9^.

**Figure 1.**
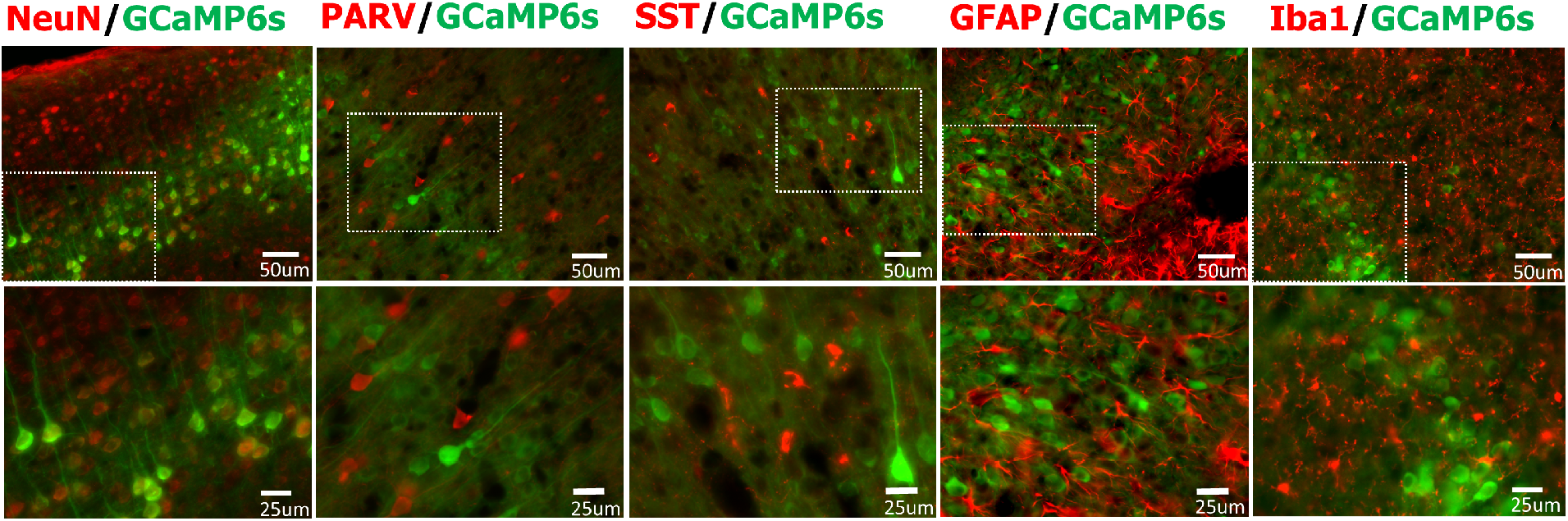
Selective neuronal expression of calcium indicator GCaMP6s. AAV9-CAG-FLEX-GCaMP6s was injected at postnatal day 1 (P1) in a representative CaMKII Cre expressing mouse. Tissue from binocular V1 was prepared at P25 and double labeled for neuronal marker (NeuN), a marker for parvalbumin+ GABAergic cells (PARV) and (SST), a marker for astrocytes (GFAP), a marker for microglia (Iba1) and GCaMP6s.

**Figure 1.**
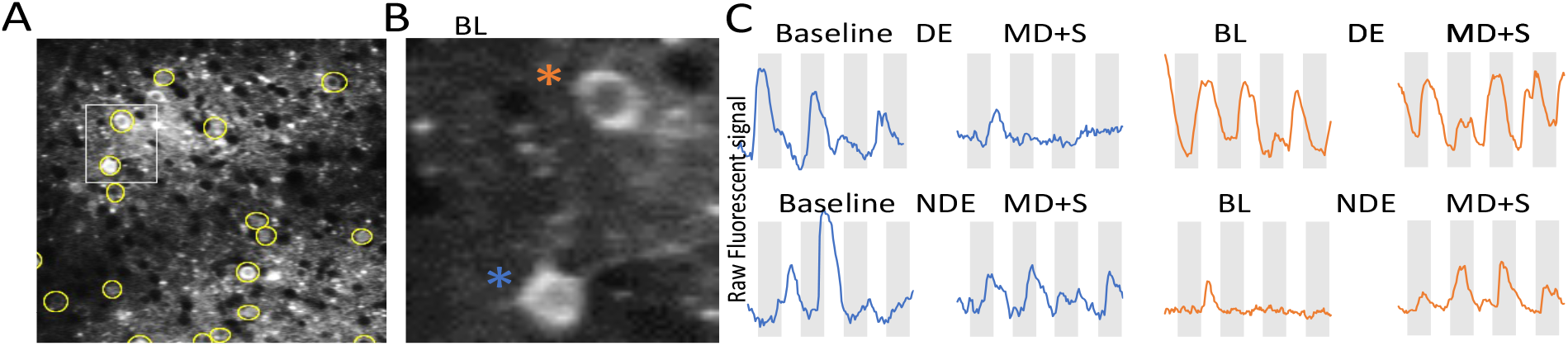
Sleep promotes bidirectional plasticity in developing visual circuits. **(A)** Yellow circles surround representative neuronal cell bodies in V1 identified by GCaMP6s 2-photon microscopy in an awake mouse during the critical period for visual plasticity. Only those neurons recorded across all experimental conditions were included for analyses. Following baseline recordings of visual responses, the mouse was subjected to 6 hours of monocular deprivation (MD) while awake + 6-hour *ad lib* sleep (+S). (**B)** 2 representative V1 neurons with different responses to MD and sleep (shown in inset box in **A**). **(C)** Calcium signal from the two neurons shown in **B.** Note that the neuron indicated by the **blue asterisk** has a weakened response to the deprived eye (DE) after sleep with minimal change in the non-deprived eye (NDE) response. In contrast, the neuron indicated by the **orange asterisk** does not show a weakened response to the DE after MD+S. Instead, the response to the NDE becomes stronger. Grey bars indicate time when gratings on a LED monitor were presented to either the DE or NDE.

We used a design adopted from previous investigations of sleep and ODP ^1, 6^. Following baseline measurements of visual responses to each eye, the eye contralateral to the measured V1 was closed and the mice were kept awake for 6 hours to induce ODP. We then determined the effects of subsequent 6 hours *ad lib* sleep or 3 hours of RSD on ODP. Undisturbed sleep after MD (MD+S) led to a shift in the ocular dominance index (ODI) which did not occur in the MD+RSD group (ODI MD+S = 0.51 +/− 0.02 (std. error) vs. baseline 0.44 +/− 0.02: paired Student’s t-test, p<0.001; MD+RSD: 0.48 +/−0.02, vs. baseline 0.47 +/− 0.02, p=0.37: paired Student’s t-test). The effects of sleep on ODP involved both a loss of response to the DE and a gain of response to the NDE (Figure 2). Averaged across neurons, both changes were enhanced after sleep compared to baseline values. These plastic changes did not occur in RSD mice (Figure 3). After RSD, changes in the NDE response were not different from baseline, and DE responses were instead enhanced.

**Figure 2.**
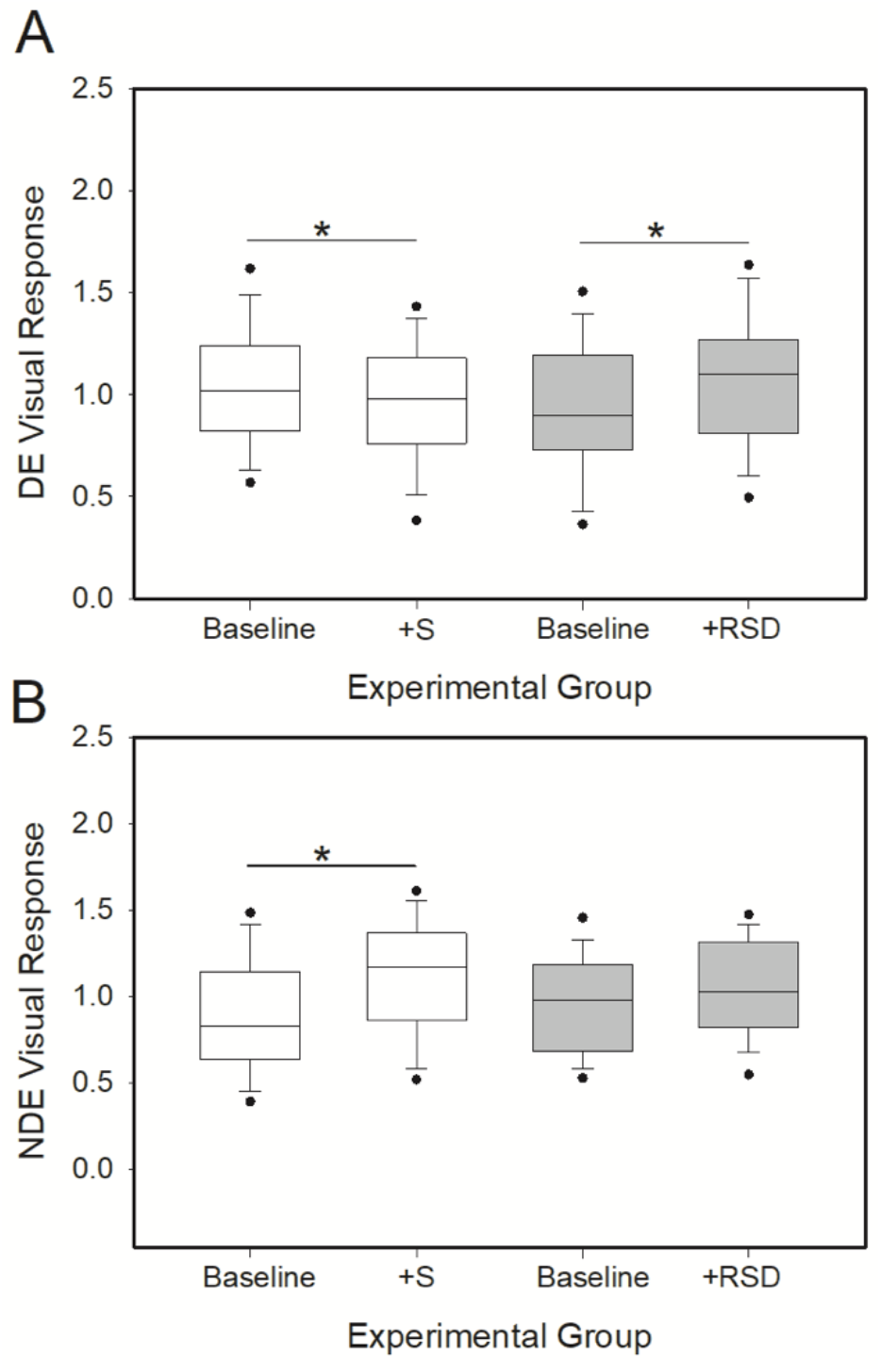
REM sleep deprivation inhibits bidirectional plasticity in developing visual circuits. Data are mean neuronal visual responses to the deprived eye (DE) and non-deprived eye (NDE) in critical period mice monocularly deprived while awake (6 hours) and then permitted *ad lib* sleep (+S), or REM sleep deprived (+RSD). (A) Relative to baseline, responses to the DE are weaker after sleep (Kruskal-Wallis, p<0.001). This weakening does not occur in RSD mice; instead, responses to the DE are enhanced (Kruskal-Wallis, p<0.05) (B) Responses to the NDE become stronger following post-MD *ad lib* sleep (Kruskal-Wallis, p<0.001); this effect is abolished by RSD (*ns*, p=0.490). 5^th^ and 95^th^ percentiles are indicated by symbols. +S= responses in sleeping group, +RSD=responses in RSD group. The number of mice and neurons per group are as follows: +S, 4, 136; +RSD, 4, 101. * Indicates significant difference between groups.

## Discussion

We investigated the role of REM sleep in a canonical model of experience-dependent plasticity (ODP). We find that undisturbed sleep after short periods of MD (6 hours) induce bidirectional plastic changes in V1 neurons as measured by 2-photon microscopy in awake mice. These results are surprising because previous studies of ODP in mice indicate that several days of MD are required for measurable changes in ODP ^10^, and sleep only promotes weakening of DE circuits ^6^. We discuss aspects of these findings below.

### Sleep-dependent ODP

The roles of different sleep states in CP plasticity are undefined. Studies in cats demonstrate that sleep enhances ODP by maintaining depression in DE circuits while strengthening responses to the NDE ^7, 11^. Cortical activity in sleep is required for both changes—as intracortical inhibition of NMDA receptors during post-MD sleep abolishes signs of circuit weakening and strengthening ^6, 7^. Our findings in mice are similar in some respects. First, like cats, we find that sleep is required for similar changes in both visual pathways as they are abolished by RSD during the first few hours of the post-MD period. However, in contrast to the present study, our earlier study in mice showed that MD followed by undisturbed sleep only produced evidence consistent with synaptic weakening ^6^.

One explanation for this apparent discrepancy is that sleep-dependent plasticity varies depending on the type of neuronal circuit ^9, 12^. For example, in adult mice motor learning followed by sleep increases dendritic spine number in motor cortex, but only on some dendritic branches ^12^. Similarly, sleep-dependent hippocampal (morphological) plasticity is highly determined by region (CA1 vs. CA3) and spatial location on a given dendrite (reviewed in ^13, 14^). Our previous study in mice examined changes in spine number and GCaMP signaling in layer 5 neuron apical dendrites ^6^. In this study, we instead examined somatic GCaMP signaling in layer 2/3 excitatory neurons. AMPA receptor trafficking (which is a proxy for changes in synaptic efficacy) varies in different V1 neurons, with more dynamic changes occurring in layer 2/3 dendritic spines compared to layer 5 apical dendritic spines ^15^. This suggests that more heterogenous plastic changes may occur in layer 2/3 neurons compared to layer 5 neurons in sleep-dependent plasticity.

Changes in ODP across the sleep-wake cycle may be explained by a Hebbian process that promotes synaptic weakening when visual experience is severely reduced and a homeostatic scaling mechanism in sleep that enhances visual responses in a manner that favors the open eye ^16^. In support of this idea, in mice the loss of response to the deprived eye is mediated by Hebbian long-term depression ^17^ (and see ^3^). However, the increased response to the non-deprived eye is governed by TNFα ^18^; a cytokine that promotes homeostatic synaptic up-scaling and is at its highest brain concentrations during sleep ^18, 19^. Although the role of sleep in homeostatic V1 plasticity is complex ^20–22^, the above findings are consistent with previous suggestions that under certain conditions, sleep may promote homeostatic synaptic upscaling ^23^.

### REM sleep mechanisms in ODP

This study supports previous work in cats and mice demonstrating that REM sleep is required for the full expression of ODP. In both species, RSD after MD reduces ODP based on a variety of measures including extracellular recording, intrinsic signal imaging, and 2-photon microscopic measures of spine morphology and intracellular calcium ^5, 6^. The underlying molecular mechanisms are not entirely known but may involve both Hebbian and non-Hebbian mechanisms. In addition to TNFα, enzymes implicated in Hebbian synaptic plasticity (ERK and mTOR) are also key mediators of ODP ^4^. In CP cats, both ERK and mTOR are activated in V1 during REM sleep^5, 24^, and inhibition of both kinases during post-MD sleep reduces ODP in a manner similar to RSD^25, 26^. Whether similar events occur in rodent ODP is unknown, but sleep deprivation in adult mice inhibits mTOR (total sleep deprivation) and ERK (RSD) in the hippocampus^27, 28^, which suggests that these mechanisms may be evolutionarily conserved.

Hebbian plasticity may also explain our peculiar observation that RSD increased responsiveness to the DE. While this may reflect slight differences in baseline values between the sleeping and RSD groups, similar paradoxical shifts in favor of the DE can occur under certain conditions. Reversible silencing of V1 combined with MD also causes a shift in favor of the DE. This has been explained as a Hebbian process where the more active pre-synaptic (NDE) inputs are punished when post-synaptic neuronal firing is out of phase ^29, 30^. Therefore, it is possible that in the absence of REM sleep, spontaneous V1 activity becomes decorrelated with pre-synaptic inputs, leading to a similar condition. While speculative, this idea is supported by the observation that cortical inhibition is enhanced during REM sleep by heightened activity of parvalbumin+ (PV) GABAergic neurons ^31^. PV neurons play essential roles in ODP by influencing the opening and closing of the CP and gating excitation and (possibly) spike-timing-dependent plasticity in pyramidal neurons following MD ^3, 32, 33^. Removing REM sleep, and these specific brain conditions, may thus disorganize the balance of cortical excitation and inhibition necessary for normal effects of MD.

## Methods

All procedures were approved by the Washington State University Institutional Animal Care and Use Committee.

### Virus injection

At P0-P3, equal numbers of male and female CaMKIIalpha-Cre mice (B6.Gg-Tg(CaMKIIalpha-Cre)T29-1Stl/J stock005359), Jackson Laboratories) were anesthetized under isoflurane 2-3% and injected bilaterally intraventricularly (stereotaxic coordinate from lambda) with a Hamilton syringe mounted to a syringe pump (0.5μl/min) with 1 μl of AAV9-CAG-Flex-GCaMP6s (titer: ~ 1×10e13 genomes/mL, Penn Vector core) and 1 μl of AAV1-CAG-tdTomato (titer: ~ 1×10e13 genomes/mL, Penn Vector core). The procedure lasted around 10 to 15 minutes ^34^. As shown previously, the former injection leads to selective expression of GCaMP6 in CaMKII+ neurons ^35^. TdTomato was used to create fiduciary marks for aligning frames. Following post-operative recovery on a heating pad, pups were returned to their home nest.

### EEG & EMG, cranial window surgery and baseplate positioning

At P18-21, mice were prepared for GCaMP6 and EEG & EMG recording as described previously^36^. Briefly, after dura removal, a 3mm diameter cranial window was positioned over binocular (right hemisphere) V1 using previously described coordinates ^37^. A baseplate that allows for head fixation for 2 photon imaging was also positioned. In the other hemisphere, three stainless-steel wire EEG (and one reference above the cerebellum) electrodes were implanted in frontal/parietal bone. Two EMG electrodes were inserted into the nuchal muscles. These electrodes were connected to a 3.1mm diameter miniature connector (Omnetics Connector Corp). The baseplate, socket and the head-restraint baseplate were secured with black (opaque) dental cement and the cranial window was protected from light with black tape.

### MD and sleep procedures

At P23-25, male mice were placed in a Pinnacle systems (http://www.pinnaclet.com/sleep-deprivation.html) sleep-recording chamber and an EEG & EMG cable and miniaturized preamplifier were connected to the socket. Mice were acclimated to the chamber and cable for 3 days prior to the beginning of an experiment (12:12h light-dark cycle, food and water *ad lib*, ambient temperature, 24-25° C). In parallel with the preceding habituation, mice were also progressively habituated to 1 hour head restraint one to two times each day for at least 5 days prior to visual testing ^6^. Following the acclimation period and beginning at zeitgeber time 0 (lights on), baseline visual responses in each eye were recorded. After this baseline measure, the eye contralateral to the hemisphere with the cranial window (left eye) was covered with a small piece of light-proof tape held by a metallic tab secured to the head bar. The mice were then kept awake with gentle handling for 6 hours to induce ODP as previously performed in cats ^1^. We then determined the effects of post-MD *ad lib* sleep or RSD on ODP in separate groups of mice. Following the above MD procedures, one group was allowed to sleep *ad lib* for 6 hours (MD+S), while the other group was REM sleep deprived (MD+RSD) for the first 3 hours post-MD followed by 3 hours *ad lib* recovery sleep. The post-MD sleep and RSD periods were conducted in dim red light to control for visual input. Between 2-photon measurements, polysomnographic signals were continuously monitored by an investigator with experience studying REM sleep in rodents (L.R.) to ensure that all mice were awake during the MD, that all mice slept during *ad lib* sleep periods, and that RSD mice were awakened during REM sleep, as previously described ^6^. This latter approach results in >90% reduction in REM sleep with non-significant changes in nonREM sleep in CP mice ^6^.

### 2-photon signal processing and visual testing

2-photon microscopy in non-anesthetized head restrained mice was performed as previously described ^36^. For visual testing, the mouse was positioned approximately 25 cm in front of a 120Hz LED monitor and each eye was presented with drifting gratings in 4 orientations (4 times per orientation at 0, 45, 90, 135 degrees, and grey screen, spatial and temporal frequency as described in ^38^, 3 seconds each separated by 6 seconds of black screen presentation). The grey screen was set at equal luminance, and zero contrast. Visual stimuli were generated using Matlab2016a and psychtoolbox. Visual responses were recorded between ~150μm and ~200μm below the window surface (visual cortical layer 2/3). A Kalman and bleaching filter (in ImageJ) was used to reduce noise and correct linear trends in the signal ^36^.

The preferred orientation (the orientation that on average produced the largest response) was determined for each neuron and used for all further calculations. Only neurons that were visually responsive in the baseline recordings (those with a mean peak grating response>mean peak response grey screen) were included in analyses. In addition, we only analyzed neurons that could be recorded across the baseline and post-MD periods. This was determined by using fiduciary marks (*e.g*., vasculature maps and TdTomato signals) to position the microscope over the same region at different points of the experiment, followed by image registration in the X and Y planes to align frames from different sets of recordings. Calcium signals obtained during visual testing were converted into ΔF/F values (ΔF/F=(F_stim_-F_spon_)F_spon_)where F_stim_= peak signal during 3 second visual stimulation (+2 seconds of subsequent black screen) and F_spon_= peak signal during the last 2 seconds of the preceding black screen presentation. This calculation ensured that slow GCaMP6s responses to a visual stimulus were also captured (Figure 1). Overall changes in ocular dominance were measured using an ocular dominance index (ODI) calculated as ODI=NDE /(DE+NDE), where DE=deprived eye, NDE=non-deprived eye. To determine specific changes in the response to each eye, each neuron’s responses were normalized to the overall mean of the baseline & post sleep or post-RSD values, respectively. The number of mice and ROIs per group were as follows: MD+S, 4, 136; MD+RSD, 4, 101.

### Statistical analysis

IBM SPSS Statistics (version 28) and Systat Sigmaplot 11 were used for statistical analysis. Data were first tested for normality. Parametric data were analyzed using Student’s t-test (paired, where applicable). Nonparametric data were assessed with the Kruskal Wallis test.

## Acknowledgements

The authors declare no conflicts of interest. This work was supported by NIH R21HD088829 (to MGF). The authors thank Dr. Wenbiao Gan for discussion and advice on 2-photon microscopy in awake mice.

